# microGWAS: a computational pipeline to perform large scale bacterial genome-wide association studies

**DOI:** 10.1101/2024.07.08.602456

**Authors:** Judit Burgaya, Bamu F. Damaris, Jenny Fiebig, Marco Galardini

**Affiliations:** Institute for Molecular Bacteriology, TWINCORE Centre for Experimental and Clinical Infection Research, a joint venture between the Hannover Medical School (MHH) and the Helmholtz Centre for Infection Research (HZI), Hannover, Germany; Cluster of Excellence RESIST (EXC 2155), Hannover Medical School (MHH), Hannover, Germany

## Abstract

Identifying genetic variants associated with bacterial phenotypes, such as virulence, host preference, and antimicrobial resistance, has great potential for a better understanding of the mechanisms involved in these traits. The availability of large collections of bacterial genomes has made genome-wide association studies (GWAS) a common approach for this purpose. The need to employ multiple software tools for data pre- and post-processing limits the application of these methods by experienced bioinformaticians. To address this issue, we have developed a pipeline to perform bacterial GWAS from a set of assemblies and annotations, with multiple phenotypes as targets. The associations are run using five sets of genetic variants: unitigs, gene presence/absence, rare variants (*i*.*e*. gene burden test), gene cluster specific k-mers, and all unitigs jointly. All variants passing the association threshold are further annotated to identify overrepresented biological processes and pathways. The results can be further augmented by generating a phylogenetic tree and by predicting the presence of antimicrobial resistance and virulence associated genes. We tested the microGWAS pipeline on a previously reported dataset on *E. coli* virulence, successfully identifying the causal variants, and providing further interpretation on the association results. The microGWAS pipeline integrates the state-of-the-art tools to perform bacterial GWAS into a single, user-friendly, and reproducible pipeline, allowing for the democratization of these analyses. The pipeline can be accessed, together with its documentation, at: https://github.com/microbial-pangenomes-lab/microGWAS.

## Introduction

Genome-wide association studies (GWAS) have gained popularity as a useful approach to identify genetic variants associated with different phenotypic traits across different organisms, including bacteria^1^. In numerous cases, variants that were statistically associated with the phenotype were confirmed as causative through follow-up laboratory experiments, demonstrating the efficacy of this approach and its value in uncovering important aspects of phenomena such as bacterial infections^2,3^. With the now general availability of bacterial genome sequencing data at low cost, GWAS has offered a promising approach to understand the genetic basis of clinically relevant bacterial traits like pathogenicity^4^, antibiotic resistance^5^ and host specificity^6^, just to mention phenotypes relevant to infection research.

Bacterial GWAS poses unique challenges compared to studies focusing on human data, as one needs to account for a mostly clonal population structure, and the presence of a large accessory genome^1,7^. This typically involves computationally intensive steps such as variant calling, population structure analysis, association testing and some post-association analyses (*e*.*g*. annotation of variants passing the association threshold, functional enrichment, generating visualizations such as Manhattan plots, etc). Despite the availability of individual computational tools to perform these analyses in bacteria, they often lack integration and user-friendliness, posing challenges for individuals with limited proficiency in bioinformatics, and with varying degrees of adherence to best practices and reproducibility^8^. There is therefore a strong need for a configurable and easy to use pipeline that implements best practices and produces reproducible results. To the best of our knowledge the available computational tools that can be classified as pipelines are CALDERA, DBGWAS and bacterialGWAS, which test for associations using k-mers from a compacted De Bruijn graph ^9–11^. These pipelines however lack a detailed downstream annotation of the variants passing the association threshold, allow only for associations to be carried out on a single type of genetic variant, have been last updated more than two years ago, and do not offer an easy way to implement reproducibility and configuration.

We have therefore developed a comprehensive microbial GWAS pipeline (microGWAS) that simplifies the entire analysis process, from raw data to the generation of interpretable results. This user-friendly pipeline uses Snakemake to ensure efficiency and reproducibility^12^. microGWAS allows for five types of association analyses with a user-defined number of phenotypes belonging to a specific dataset, and includes the estimation of heritability, functional enrichment analysis of the output associations, and common diagnostic and interpretative visualizations. This brings together all the necessary steps to perform a GWAS analysis for a variety of bacterial datasets, facilitating researchers with limited computational experience to leverage these methods for their bacterial genomics studies and improving their reproducibility. The pipeline, together with extensive documentation and a small test dataset, can be accessed at: https://github.com/microbial-pangenomes-lab/microGWAS.

## Methods

### Overview of the microGWAS pipeline

The microGWAS pipeline is implemented using Snakemake, a workflow management system (**Figure 1**). The pipeline runs on a set of assembled genomes and one or multiple phenotypes. Associations are run using five different sets of genetic variants: unitigs, gene presence/absence, rare variants (*i*.*e*. gene burden test), gene cluster specific k-mers, and all unitigs combined (*i*.*e*. whole genome machine learning model). Downstream analyses are carried out for each set, providing annotated hits and functional enrichments as outputs. Additionally, the pipeline estimates the heritability of each phenotype using the lineage information and the unitigs. The pipeline can also be used to generate a core genome phylogenetic tree of all samples, which can be visualized together with the distribution of the input phenotypes and number of GWAS hits using Microreact^13^. Finally, antimicrobial resistance (ARGs) and virulence associated genes (VAGs) can be predicted and categorized into functionally relevant groups. The pipeline is available as a code repository on GitHub (https://github.com/microbial-pangenomes-lab/microGWAS), for which we also provide extensive documentation: https://microgwas.readthedocs.io. In the following sections, we provide the versions of the tools used as part of the pipeline during its development and testing.

**Figure 1.**
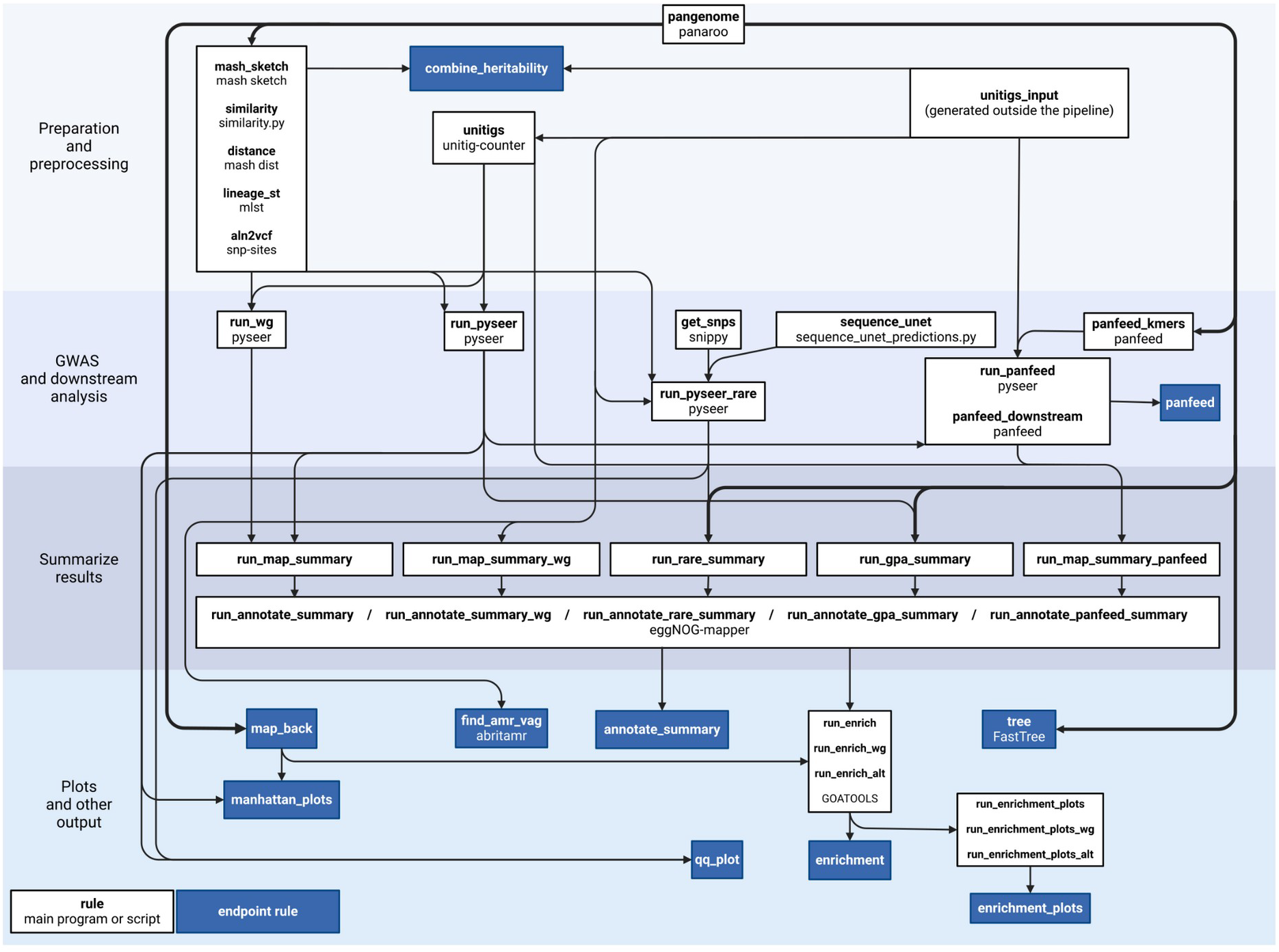
Overview of the microGWAS pipeline. Simplified directed acyclic graph (DAG) of the pipeline processing steps. The name of each Snakemake rule is reported in bold, and the name of the underlying tool or script used by each rule is specified below the rule name.

### Input data pre-processing

The pipeline requires a table encoding the target phenotype(s), any potential covariant, and the path of each sample’s nucleotide sequences (in FASTA format) and annotations (in GFF3 format). From this file, the user can use a convenience bootstrapping script `bootstrap.sh`) to prepare the actual inputs of the pipeline. The script will also download the reference genomes used for the annotation of hits and for the rare variants analysis, using ncbi-genome-download v0.3.3^14^. Multi-locus sequence typing is performed using mlst v2.16^15,16^. Genes related to virulence and antimicrobial resistance are identified by running abritamr v1.0.14^17^ on the nucleotide sequences of the isolates.

### Generation of genetic variants

Associations are run on four different genetic variants: unitigs, gene presence-absence, gene cluster k-mers and rare variants.

### Unitigs

A de Bruijn graph from the nucleotide sequence of all input genomes is generated and unitigs and their presence/absence patterns are extracted across all samples using unitig-counter v1.1.0^10^. Each unitig with allele frequency < 1 encodes for a genetic variant, which can be a large one such as a gene transferred horizontally, as well as short variants such as single nucleotide polymorphisms (SNPs).

### Gene presence-absence

A pangenome of all input isolates plus the chosen references is computed with panaroo v1.3.0^18^, using the “--clean-mode strict” argument, generating a presence/absence matrix for all identified orthologous genes.

### Gene cluster k-mers

panfeed v1.6.1^19^ is used to extract k-mers and their presence-absence patterns from each gene cluster identified by panaroo separately. Each k-mer is directly linked to its source gene, improving interpretation and showing the local context of the associated variants.

### Rare variants

Short variants (SNPs and InDels) with frequency < 5% and a predicted deleterious impact are retained to carry out associations through gene burden testing. Each input genome is mapped against the reference chosen by the user using snippy v4.6.0^20^, and then merged and normalized using bcftools v1.13^21^. Only frameshift, nonsense, and non-synonymous variants are kept for the generation of the final VCF file. Non-synonymous variants are further filtered so that only those that are predicted to be deleterious are kept; Sequence UNET v1.0.6^22^ is used to score each non-synonymous variant, with a probability threshold of 0.5.

### Estimation of heritability

Narrow-sense heritability is estimated for each of the target phenotypes, using two different covariance matrices; one built from the lineage of each strain and another using a kinship matrix built from the unitigs presence and absence matrix derived from the input genomes. Limix v3.0.4^23^ is used for the estimation, assuming normal errors for the point estimate, and computing 95% confidence intervals using the ALBI package^24^.

### Association analyses

The distance between each pair of samples is computed by using mash v2.2.2^25^ with a sketch size of 10,000; the resulting distance square matrix is utilized to compute associations between lineages (STs) and each target variable, using pyseer v1.3.6 ^26^. The association between each unitig with minimum allele frequency (MAF) > 1% and each phenotype is tested employing a linear-mixed model (LMM) as implemented in pyseer. We determine an appropriate significance threshold by counting the number of unique unitigs presence/absence patterns tested, which reduces the risk of excessively deflating association p-values. The same threshold derived from the single locus unitigs association is used for the other four associations for consistency. Users should consider employing a more stringent p-value threshold when testing a large (*e*.*g*. > 5) number of phenotypes utilizing this pipeline. The unitigs are further filtered to reduce the number of spurious associations: unitigs are excluded if they are shorter than 30bp, if they are mapped to multiple locations in each individual genome, if they map to less than 9 samples, and if they are mapped to more than 10 different genes across all samples.

In order to run associations between gene presence-absence and the phenotype, a kinship matrix is required. This can be derived from a core genome alignment, obtained through individual alignments for each nucleotide coding sequence belonging to core genes (*i*.*e*. with frequency >= 95%). Subsequently, snp-sites v2.5.1^27^ and bcftools v1.13^21^ are employed to convert the full core genome alignment to a VCF file containing the variant sites with MAF > 1%, which is then utilized file to derive the kinship matrix via a python script. Gene cluster specific k-mers are also used to test for associations with the phenotype, as implemented in panfeed v1.6.1^19^. To test for rare variants, we focus on non-synonymous SNPs against a reference chosen by the user, with a frequency < 5%, and predicted to have a deleterious effect. Finally, an association employing the full set of unitigs is tested by training a linear model with elastic net regularization using the presence/absence patterns of all unitigs. Two models are trained using an alpha parameter with a value of 1 and 0.1, which are equivalent to a lasso and ridge model, respectively.

### Annotation of threshold-passing variants

For each set of genetic variants, a number of downstream analyses is run to provide annotated results and functional enrichments. The unitigs passing the significance threshold are mapped back to all input genomes employing bwa v0.7.17^28^ and bedtools v2.31.1^29,30^, using the output of panaroo to assign each unitig to a gene cluster. Gene families with at least one unitig mapping to them are further annotated by taking a representative protein sequence from the genomes encoding each gene family, giving priority to the reference genomes indicated by the user, and using them as an input for eggnog-mapper v2.1.3^31^. The derived annotations include clusters of orthologous gene (COG) categories^32^; gene ontology (GO) terms, and mapping to KEGG entries^33–35^. We test an enrichment for each of these annotation systems using the associated gene families as the foreground and the annotation for the chosen reference genome as the background, running a Fisher’s exact test for each annotation item, and using an FDR-corrected p-value threshold^36^ of 0.05 to indicate annotation items enriched in the associated gene families. For GO terms, we utilize the enrichment tests implemented in goatools v1.2.3^37^. Plots visualizing these enrichment results are further generated. Additionally, Manhattan plots based on the mapping of all tested unitigs on the chosen reference genome are created for unitig association results, as well as diagnostic QQ plots for all single locus associations.

## Results

### Faster development using a reduced test dataset

We have included a small proof-of-concept dataset in the pipeline in order to quickly and reliably test the pipeline. This ensures that we can apply a continuous integration testing paradigm into maintaining and further developing the pipeline, which results in a more sustainable software and facilitates code additions from the community. The test can be run on a laptop with 8 cores and at least ∼10Gb of RAM in a few hours. We have created the test dataset from a previously published dataset from a mouse model of bloodstream infection^3^. To run the test, the user only needs to create a symbolic link to the eggnog-mapper database and execute the `test/run_test.sh` bash script. This script will prepare the input files, run the bootstrapping script, and run Snakemake, initially in “dry” mode printing the commands that have to be performed and then actually performing the run.

### Validation of the pipeline on an *E. coli* virulence dataset

We validated the outputs of the microGWAS pipeline on a previously published collection of 370 commensal, pathogenic and environmental strains representative of the *Escherichia* genus phylogenetic diversity^3^. In this study, a mouse model of sepsis was used to characterize the virulence phenotype of the strains, for which genomic sequences were also available. The GWAS was performed using the linear mixed model implemented in pyseer and correcting for population structure. The GWAS performed in the original study found significant variants (*i*.*e*. unitigs) related to three iron-uptake systems: the high-pathogenicity island (HPI), aerobactin, and the *sitABCD* operon. Variants within the core genes were also identified, including *zinT, mtfA* or *shiA*, most of which are encoded around the HPI. These variants were experimentally validated, showing virulence attenuation in strains with deletions in the *irp2* HPI gene.

We could replicate the findings from the original study using the microGWAS pipeline, and in addition, we automatically generated interpretative visualizations on the associated variants and functional enrichments (**Figure 2** and **Figure 3**). The pipeline also generated easily parsable outputs that can be used to generate other visualizations in a straightforward manner to provide further information on the association analysis, such as zoomed in regions for the Manhattan plots (**Figure 2C**), and volcano plots (**Figure 3A**). We found associations with all three iron capture systems that were identified in the previous study, using unitigs, gene presence/absence patterns, and gene-cluster specific k-mers. The lasso whole-genome model had a R^2^ of 0.48. Among the genes with non-zero coefficients, we identified additional variants: *iroN* and *iroC*, related to iron acquisition and virulence. These variants also interact with the HPI^38^, *hha*, a haemolysin expression modulating protein^39^, and *norW*, which is involved in anaerobic respiration, by using NADH to detoxify nitric oxide^40^. The ridge model had instead a higher R^2^ (0.71) and we found 888 genes with a significant model coefficient, which can be expected given its wider regularization during model training. All association modalities, except the gene burden test, validated the earlier results. We note that in order to obtain these results, we only had to prepare a table encoding the virulence phenotype and the path to the annotated genome assemblies, and to edit the pipeline configuration file. This demonstrates the potential of this pipeline to facilitate bacterial GWAS by removing the barriers imposed by the need to orchestrate multiple bioinformatic tools and their individual idiosyncrasies.

**Figure 2.**
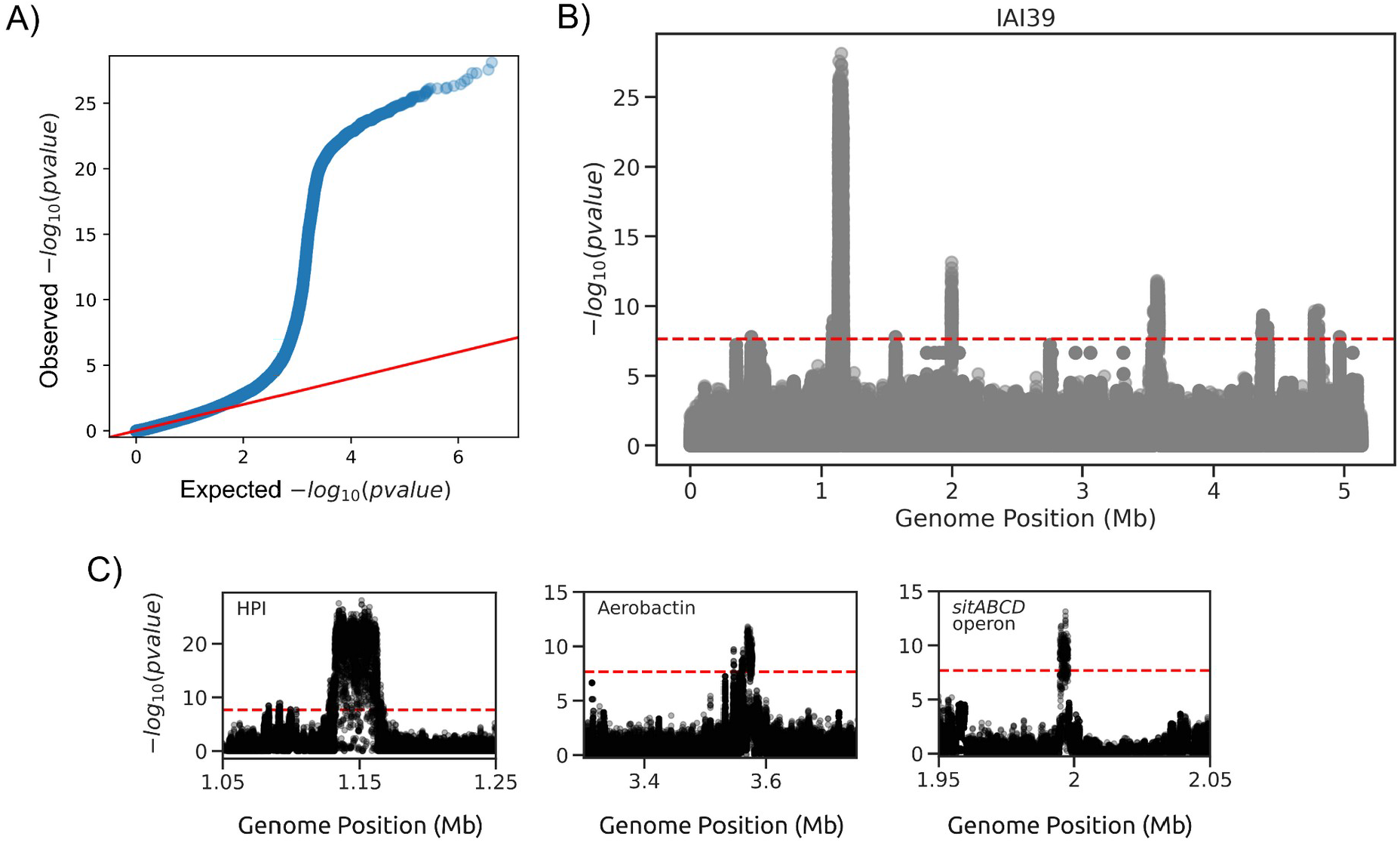
Representative visualizations on the associated variants based on unitigs. Panels A and B have been generated directly by the pipeline, while panel C has been generated directly from output files. A) QQ plot showing the distribution of observed p-values with the expected distribution under the null hypothesis. B) Manhattan plot of the associated variants, using strain IAI39 as a reference. The significance threshold is indicated by the red dashed line. C) Zoom-in on the associated areas of the Manhattan plot for the HPI, aerobactin and *sitABCD* operon regions.

**Figure 3.**
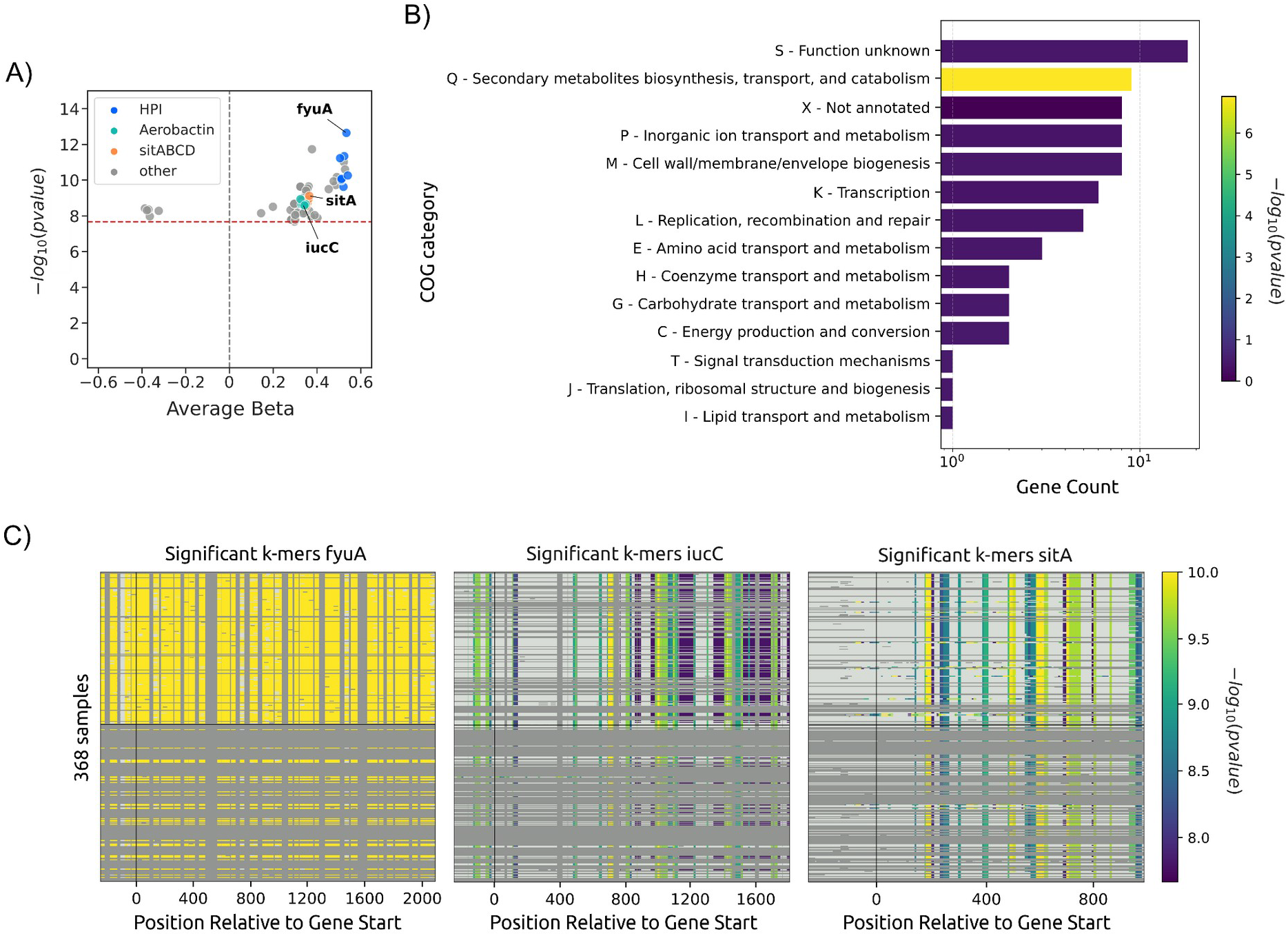
Representative visualizations on the associated variants and functional enrichment based on unitigs. Panel A has been generated directly from output files, while panels B and C have been generated directly by the pipeline. A) Volcano plot of the associated variants. Genes associated with the high-pathogenicity island (HPI), the aerobactin, and the *sitABCD* operon are highlighted. The significance threshold is indicated by the red dashed line. B) Enrichment analysis of the associated unitigs for the different COG categories. The x-axis shows the number of gene hits belonging to each category. The bars are coloured according to -log_10_ of the enrichment corrected p-value. C) Association plots for gene cluster specific k-mers. Each panel shows the significant k-mers for the *fyuA, iucC*, and *sitA* genes. The y-axis represents each isolate, and the x-axis the k-mer positions relative to the gene start codon for each strain. The colors correspond to the -log _10_ of the association p-value. The dark gray regions imply that the isolates do not encode for the k-mers, while the light gray regions represent k-mers under the association threshold.

## Discussion

Bacterial genome associations studies have emerged with a considerable time lag with respect to the field of human genetics. The reasons can be traced back to a lack of sufficiently large datasets, but also to the peculiar genetic architecture of bacterial populations, which required the development of dedicated tools capable of properly correcting the association analyses^8^. These bacterial GWAS tools in turn rely on a large ecosystem of software tools designed to preprocess the input genomes, extract genetic variants, define the so-called pangenome, as well as downstream analyses such as annotating gene hits and run functional enrichments on them. This requires the user to be familiar with each individual tool and their requirements; as a result, only experienced bioinformaticians are able to carry out these analyses with relative ease. This complexity can also lead to a lack of consistency in adhering with best practices in data preprocessing and in the actual associations, with the risk of users running poorly controlled analyses and reporting spurious results. We therefore saw the need for a user-friendly solution that abstracts away these hurdles and democratizes bacterial GWAS for less experienced bioinformaticians.

We have developed a comprehensive pipeline for conducting bacterial GWAS. This pipeline simplifies the entire GWAS process by taking as inputs bacterial annotated genome assemblies, together with a table specifying the phenotype(s) that are targeted by the association analysis, and produces interpretable results for the user. The pipeline utilizes four different types of genetic variants (unitigs, gene presence/absence, rare variants, and gene cluster specific k-mers) and associates their presence with the user’s given phenotype(s). It also maps significantly associated variants against a reference genome and generates diagnostic visualizations. Additionally, we estimate heritability for the target variables using the genetic variants and the sequence types. We have also enabled multiple enrichment analysis of the associated gene families, employing a user chosen reference genome as a background for the enrichment tests. We have provided detailed online documentation to help the users set up and run the pipeline, and navigate the outputs. Finally, we have included a small test dataset that can be run in a standard laptop to provide a continuous integration environment for bug fixes and further development. This is a rare occurrence in the field and improves the future sustainability of the pipeline, by providing an entry point for junior bioinformaticians wishing to extend the pipeline.

We have demonstrated the validity of the pipeline by reproducing the findings of an earlier study. This required the preparation of a single, simple input file, and the editing of the pipeline configuration file to indicate the name of the target phenotype and the reference genomes to be used. We proceeded to follow the instructions provided in the online documentation and simply waited for the pipeline to complete. The output files could then be parsed in a straightforward manner, allowing the same conclusions to be reached about the gene clusters associated with virulence in the *E. coli* BSI mouse model. This ease of use will hopefully be appreciated by the community, leading to more studies linking bacterial genetic variability to relevant phenotypes.

## Author contributions

MG wrote the first draft of the pipeline. All authors contributed code to the final version of the pipeline. JB, BFD, and MG wrote the online documentation. JB, BFD, and JF thoroughly tested the pipeline on a number of diverse datasets and reported/fixed bugs. JB, BFD, and JF prepared the figures. All authors wrote and edited the manuscript.

## Acknowledgments

JB, BFD and MG were supported by the Deutsche Forschungsgemeinschaft (DFG, German Research Foundation) under Germany’s Excellence Strategy - EXC 2155 - project number 390874280. JB and BFD were further supported by the Hannover Biomedical Research School (HBRS), and the Center for Infection Biology (ZIB). BFD was further supported by the Graduate School Scholarship Program (GSSP) from the German Academic Exchange Service (DAAD).

